# Physically interacting beta-delta pairs in the regenerating pancreas revealed by single-cell sequencing

**DOI:** 10.1101/2021.02.22.432216

**Authors:** Eran Yanowski, Nancy-Sarah Yacovzada, Eyal David, Amir Giladi, Diego Jaitin, Lydia Farack, Adi Egozi, Danny Ben-Zvi, Shalev Itzkovitz, Ido Amit, Eran Hornstein

**Author notes:** Correspondence should be addressed to: Eran Hornstein, PhD, MD., Department of Molecular Genetics, Weizmann Institute of Science, 234 Herzl St., 76100 Rehovot, Israel.

## Abstract

The endocrine pancreas is able to regenerate in response to insult, including by driving beta-cells into the cell division cycle. Until recently, communication between neighboring cells in islets of Langerhans was overlooked by single-cell genomic technologies, which require rigorous tissue dissociation into single cells. Here, we utilize sorting of physically interacting cells (PICs) with single-cell RNA-sequencing to systematically map cellular interactions in the regenerating endocrine pancreas. The cellular landscape of the regenerated pancreas features regeneration-associated endocrine populations.

We explore the unexpected heterogeneity of beta-cells in regeneration, including an interaction-specific program between paired beta and delta-cells. Our analysis suggests that the particular cluster of beta-cells that pair with delta-cells benefits from stress protection, implying that the interaction between beta and delta-cells safeguards against regeneration-associated challenges.

## Introduction

Islets of Langerhans of the endocrine pancreas contain five main cell types that secrete hormones, which are critical for regulating multiple aspects of metabolism. Somatostatin acts in a paracrine manner to inhibit other endocrine cells, including insulin secretion from beta-cells and the secretion of the counterregulatory hormone, glucagon, from alpha-cells (Strowski et al., 2000). Additional endocrine cell types that express somatostatin receptors, such as pancreatic-polypeptide cells, might be under paracrine control of somatostatin (Ludvigsen et al., 2004).

Better understanding of the cellular and molecular mechanisms underlying pancreas regenerative ability, may contribute to developing therapeutic treatment for diabetes. Accordingly, several experimental paradigms were established for the study of pancreas regeneration in rodents including injury models (Bonner-Weir et al., 1993), pancreatic duct ligation (Xu et al., 2008), and chemical ablation of islet cells (Fernandes et al., 1997). Such paradigms have been often used in concert with lineage tracing to decipher the origin of regenerating endocrine cells. Thus, new insulin-producing cells arise primarily from existing beta-cells (Dor et al., 2004), but in some conditions also from conversion of pancreatic alpha-or delta-cells (Chera et al., 2014).

Partial pancreatectomy (Ppx) is a well-characterized experimental model for initiating pancreas regeneration. Partial pancreatectomy, induces rapid endocrine and exocrine tissue regeneration, in young rodents that peaks after a week. Animals re-gain euglycemia at about 4 weeks (Jonas et al., 2001; Laybutt et al., 2002; Lee et al., 2006;Peshavaria et al., 2006). However, the regeneration capacity declines sharply with age (Rankin and Kushner, 2009) and is absent in adult humans (Menge et al., 2008; Zhou and Melton, 2018).

Much is known about beta-cell mass replenishment. New beta cells are primarily descendants of mature beta-cells that re-enter the cell division cycle (Dor et al., 2004; Lee et al., 2006). In addition, auxiliary pathways, commit beta-cells from other trajectories (Tritschler et al., 2017).

Delta-cells may contribute to regeneration of the endocrine pancreas in diverse ways, including via paracrine somatostatin activity (Alberti et al., 1973; Fagan et al., 1998) or physical cellular interaction with beta-cells (Arrojo et al., 2019). However, delta cell involvement in regeneration was largely overlooked.

Single-cell transcriptomics technology enables a new and thorough understanding of pancreas tissue dynamics and of cellular heterogeneity (Wang and Kaestner, 2019). Beta cells display transcriptomic profiles, associated with proliferation (Wang et al., 2016) and cellular stress (Baron et al., 2016; Muraro et al., 2016) and may further reveal the molecular signature of type 2 diabetes (Segerstolpe et al., 2016), post-pancreatectomy (Tatsuoka et al., 2020) and the aging organ (Xin et al., 2016).

In the current study we demonstrate that delta-cells are Sox9 descendants that expand in the regenerating endocrine pancreas. We characterize regeneration-associated beta cell heterogeneity at a single-cell resolution, demonstrating molecular patterns that are consistent with activation of stress programs or of the cell division cycle in beta-cells. Furthermore, we identify a cluster of beta-delta cell pairs that display unique transcriptional profiles. Better understating of crosstalk between endocrine cells in regenerating islets of Langerhans may contribute to our understanding of how beta-cells cope with stress and to development of therapy for diabetes.

## Results

### Islet Sox9-descendant cells are predominantly delta-cells

The transcription factor Sox9 plays critical roles in the development of the embryonic pancreas, whereas it is predominantly expressed in the exocrine compartment of the adult organ. Accordingly, Sox9-descendant cells contribute to the ductal and acinar system. In analysis of Sox9-CreER:tdTomato allele (Soeda et al., 2010), we characterized the lineage signal 30 days after labeling and observed cells within islets of Langerhans, in addition to ductal and acinar cells (Figure 1A). The islet cells that are linked to the Sox9 lineage primarily express the hormone somatostatin (Sst), which is typical of endocrine delta-cells (Figure 1B-D). Quantification of the signal demonstrated that 78% of the tdTomato^+^ cells are Sst^+^, whereas 9% co-express Insulin and 13% were neither delta nor beta-cells (Figure 1E, 1905 cells counted from 32 islets, obtained from 5 animals). Accordingly, 18% of the delta-cells were tdTomato^+^, in contrast to only 0.4% of the beta cell population (54 tdTomato^+^, Sst^+^ /295 delta-cells; 6 tdTomato^+^, Ins^+^ /1595 beta cells Figure 1F). Therefore, a relatively specific activation of the endogenous Sox9 promoter is evident in adult delta-cells or their ancestors, in the endocrine pancreas.

**Figure1.**
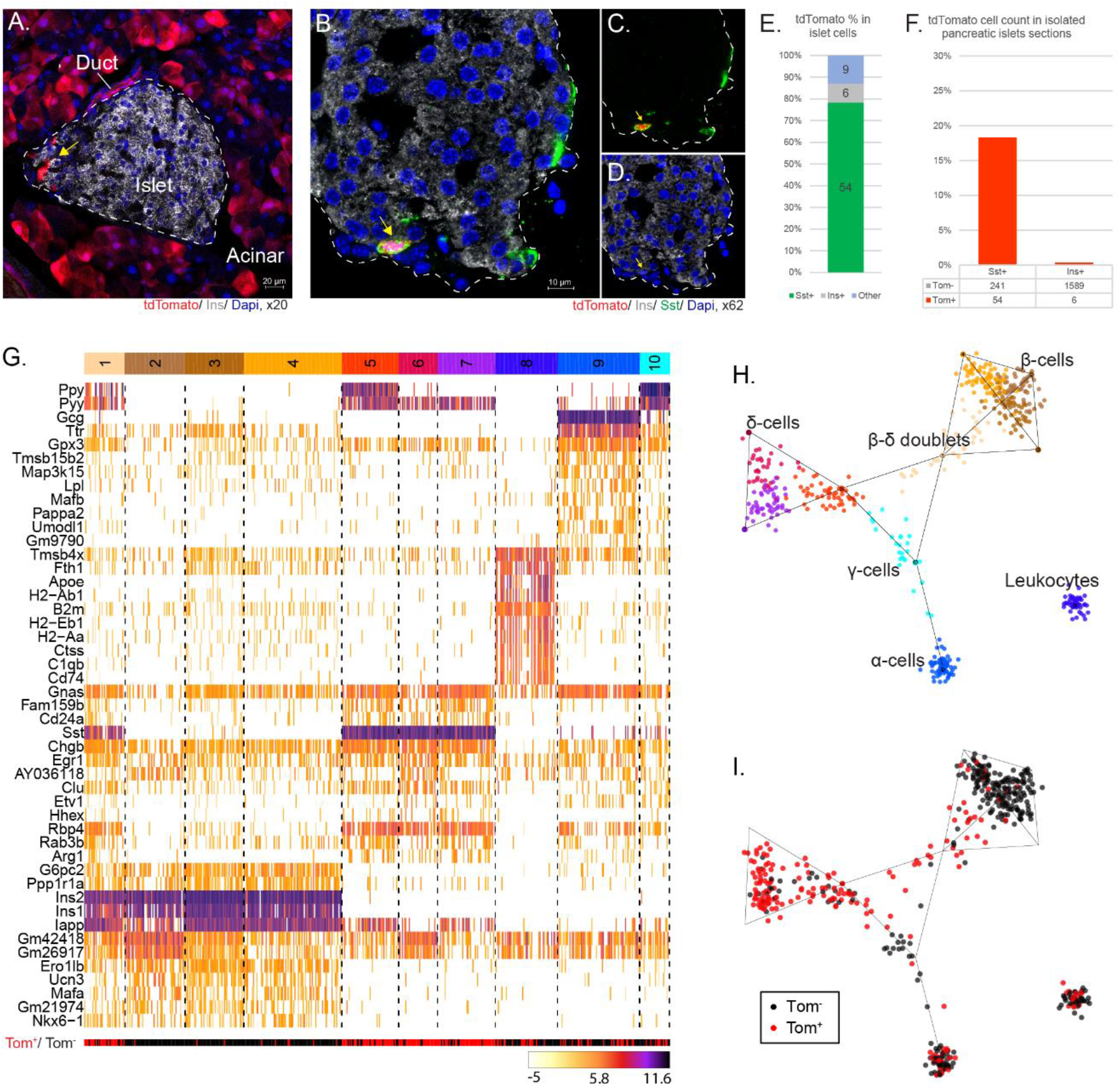
Delta-cells are Sox9 descendants. Confocal micrographs demonstrating that **(A)** Sox9 lineage tracing marks duct and acinar cells and some cells at the periphery of Islets of Langerhans (Marked by yellow arrow). Islet border is marked by white dashed line. **(B)** Sst+ delta-cells are Sox-9 descendant cells. tdTomato signal (red, yellow arrow), insulin (white) Sst (green) and nuclei (blue) co-staining. **(C)**, compared to Insulin immunostaining and DAPI **(D)**. The Islet border is marked by a white dashed line. **(E)** Quantification of Sox-9 descendant fractional enrichment in endocrine populations: delta-cells (78%), beta-cells (9%) or other islet cells (13%) (.**(F)** 20% of Sst+ cells (54 of 295 cells) and ~0.4% of Ins+ cells (6 of 1595 cells) are Sox9-descendants (tdTomato+). **(G)** Heatmap of unsupervised clustering of single endocrine cell transcriptome from 590 dissociated cells from 6 pancreata of Sox9 lineage-tracer mice. Analysis reveals that Ppy, Pyy and Sst mRNAs are enriched in lineage traced tdTomato+ cells (bottom red / black bar). Ten key cell type clusters coded by a spectral bar. **(H)** Dimensional reduction of transcriptomic data, with cell-types spectral color, corresponding to Fig. 1G. **(I)** Color coded SOX9-descendants (red) or cells unlabeled with tdTomato during the course of the experiment, superimposed on the dimensional reduction map, highlights the enrichment of Sox9-descendant delta-cells.

To uncover the nature Sox9-descendant cells, pancreata of 12 weeks old mice were dissociated and MARS-seq was performed for both tdTomato^+^ and tdTomato^-^ cells (Baran et al., 2019; Jaitin et al., 2014). Transcriptional profile analysis identified 5 distinct cellular populations, namely, beta-, delta-, gamma- and alpha-cells and circulating leukocytes (Figure 1E, F). In addition, we identified a cluster of cells holding transcriptional signature that is typical of pairs of both beta and delta-cells. These were therefore regarded as beta-delta doublet cells (cluster #1, Figure 1E). Notably, Sst mRNAs is enriched in lineage tdTomato+ cells. This analysis orthogonally demonstrated that tdTomato-labelled Sox9-descendants are indeed delta-cells (Figure 1G - I).

### Delta-cell expansion in pancreas regeneration

The adult pancreas is capable of regenerating in response to injury. However, the regeneration process is only partially characterized. Therefore, we aimed to investigate the cellular dynamics of pancreas regeneration by single cell sequencing (Figure 2A). Partial (subtotal) pancreatectomy (Ppx), was performed on 12 week-old Sox9-CreER;tdTomato males. Excision of ~70% of the pancreas was controlled by a cohort of sham operated littermates. Incomplete normalization of blood glucose levels was monitored over 4 weeks after the surgical procedure (Supplementary Figure S1A). A significant increase in the number of tdTomato^+^ cells, was observed in regenerating pancreata, 4 weeks after pancreatectomy, relative to sham operated mice (Figures 2, B-D). We have demonstrated, by single molecule fluorescent *in situ* hybridization (smFISH), an increase in the number of cells expressing Sst, 4 weeks after Ppx, relative to sham controls and to an earlier time point, 1 week after surgery (Figures 2, E-G). Together, the delta-cell population expands during regeneration.

**Figure 2.**
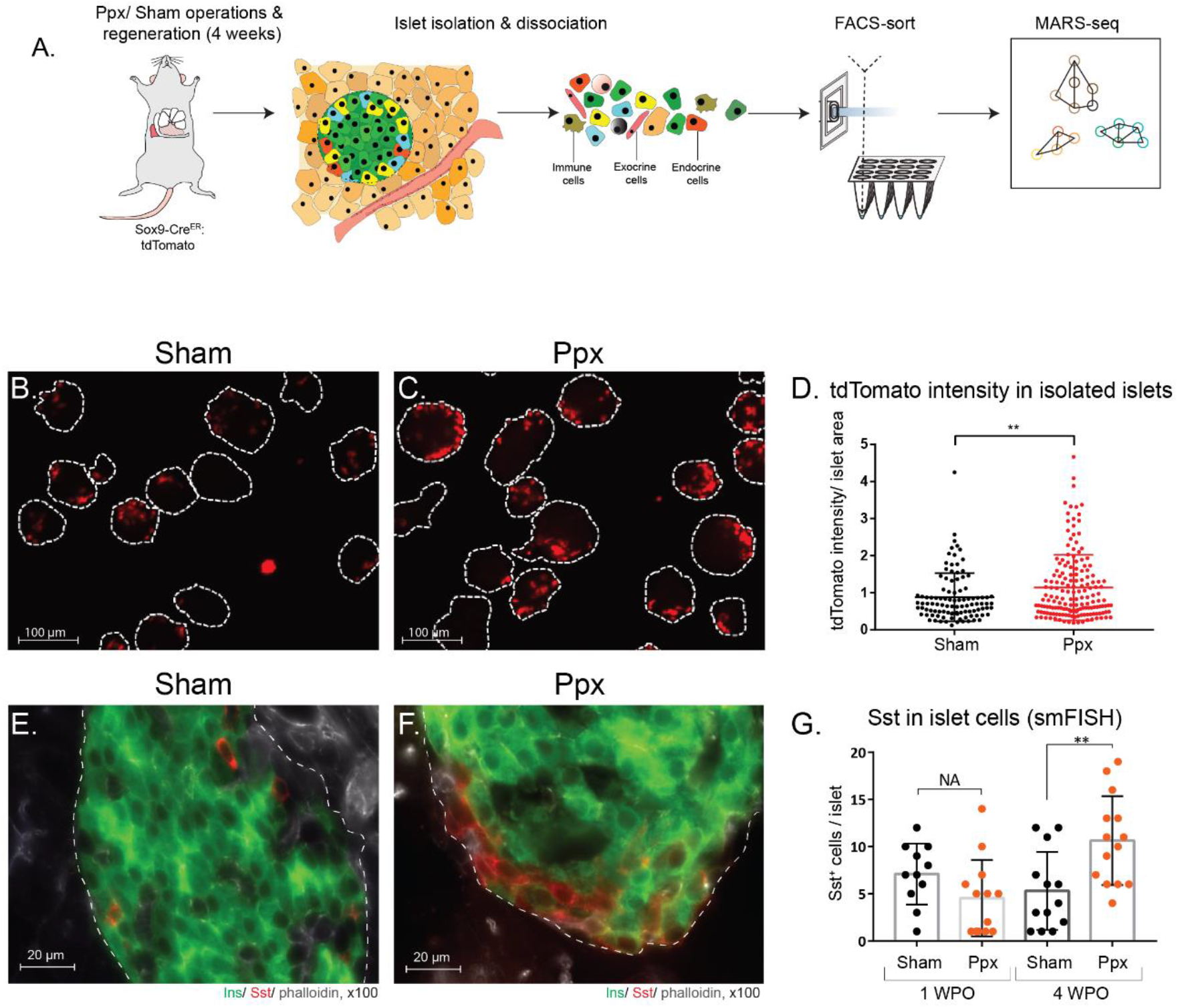
Delta-cells expand following partial pancreatectomy. **(A)** Diagram of experimental setup for single cell RNA sequencing of endocrine cells from the regenerating pancreas. Confocal micrographs of Sox9-descendant (tdTomato^+^) cells enrichment in islets of sham operated mice **(B)**, relative to after pancreatectomy **(C)**. White line - islet border. **(D)** Quantification of tdTomato intensity in isolated islets, normalized to islet area. tdTomato signal in sham: 0.88 ± 0.06 (pixels, fluorescence intensity mean value), n=102 and PPX: 1.14± 0.07 (pixels, fluorescence intensity mean value), n=165. Two-tailed T-test p = 0.0059. Confocal micrographs demonstrating delta cell hyperplasia (red) in **(E)** sham-operated mice and **(F)** regenerating islets, 4-weeks post pancreatectomy. **(G**) Bar graph quantification of Sst+ cells per islet in sham or Ppx operated mice, following one-or four-weeks post operations (WPO). One-way ANOVA, p-value = 0.0016. 355 Sst+ cells recorded in 51 islets from the 4 experimental groups.

### Single-cell characterization of the regenerating endocrine pancreas

Next, we dissociated islets from regenerating pancreata, annotated tdTomato by flow cytometry, and performed single-cell RNA sequencing. Four endocrine cellular populations, (alpha, beta, gamma and delta-cells), leukocytes, endothelial and mesenchymal cells were detected (Figure 3A, B), consistent with the profiling at basal conditions in Figure 1.

**Figure 3.**
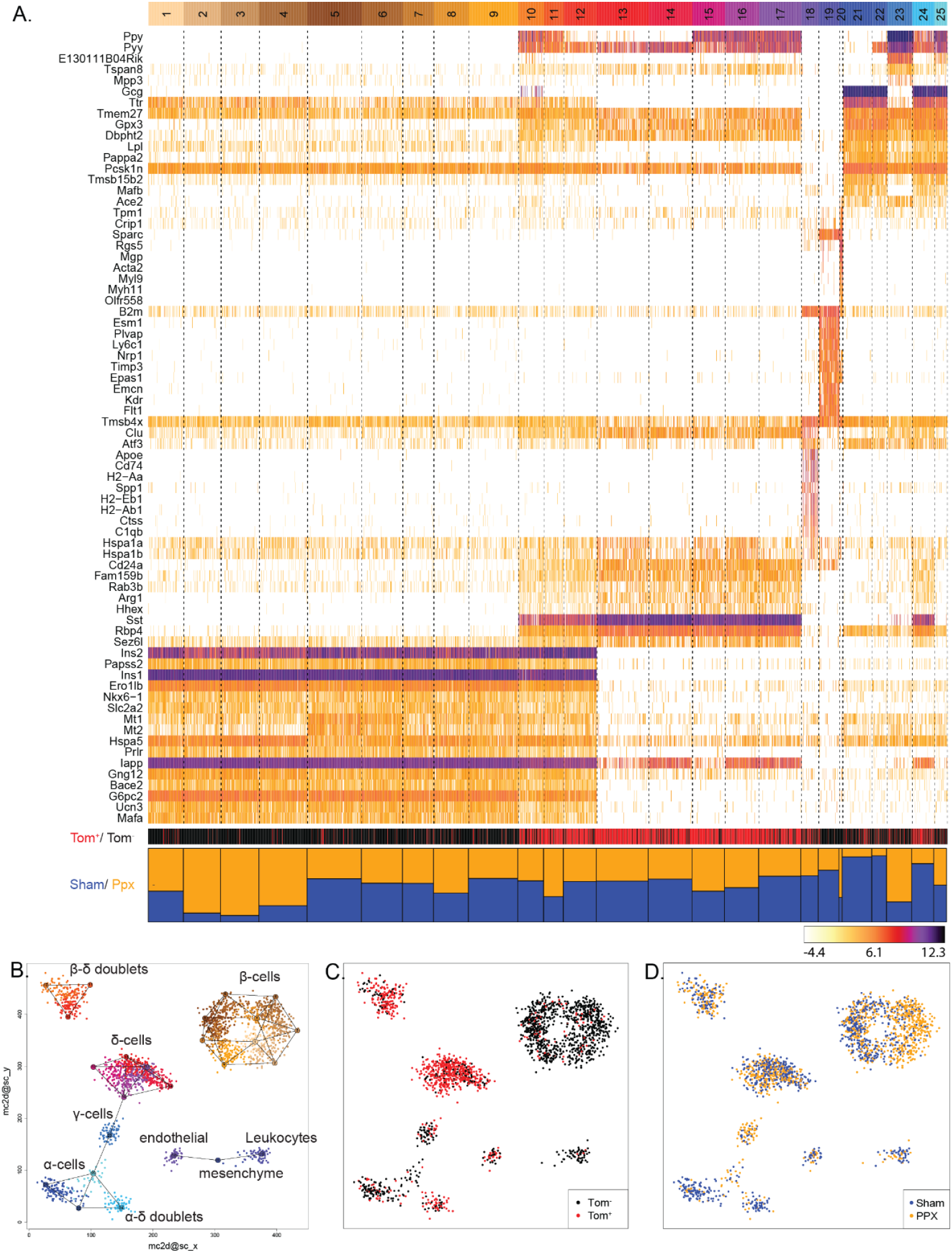
Single-cell map of the regenerating islet. **(A)** Heatmap of clustered islet cells RNA-sequencing data from 2300 cells collected from Sox9 lineage-traced 6 pancreata, 4 weeks after partial pancreatectomy. Sham-operated mice and pancreata without lineage tracing included in the same analysis. Meta-cells (columns) and maximally enriched gene markers (rows) analyzed as in (Baran et al., 2019). Color-bars indicate cell partitioning (Sham (blue) or ppx (orange)) and the presence of tdTomato (Tom, red, marker of lineage traced Sox9 descendants). Two-dimensional projection of Meta-cells and inferred cell-types **(B)**, lineage traced Sox9 descendants (tdTomato, red, **C**) and by source from sham operated or regenerating pancreata **(D)**.

The enrichment of delta-cells in this study, via Sox9 tracing, enabled analysis of the delta-cell population at unmet sensitivity (Figure 3C), while highlighting the source of cells according to their experimental groups (sham/ ppx) revealed relative depletion of alpha cells in ppx samples (only 20% originate from ppx samples), enrichment of gamma-cells (~80% originate from ppx samples) and regeneration-driven heterogeneity within the beta-cell population (Figure 3D).

### Heterogeneity of beta-cells during recovery period after pancreatectomy

In-depth analysis of beta-cell mRNA profiles revealed acquired heterogeneity in response to regeneration. In addition to a relatively-stable beta-cells that do not transcriptionally change in response to regeneration (Figure 4A), other beta cells feature elevated levels of stress-associated transcripts (Fkbp11, Dapl1, Creld2, Sdf2l1, Pdia4 & Derl3. Figure 4B), whereas other beta cells can be denoted by cell-cycle-associated transcripts (e.g., Top2a, Ccna2, Ube2c, Cenpf & Nusap1. Figure 4C). The three signatures feature tonic expression of Ins1, but can be further distinguished by discrete levels of ins2 expression (Figure 4D). Thus, beta cells with stress-associated transcripts also possess relatively low levels of Ins2 mRNA (Figure 4D, E). In summary, beta-cell heterogeneity increases in the regenerating pancreas denoting three primary transcriptional states: stress-associated, cell-cycle-associated or basal state.

**Figure 4.**
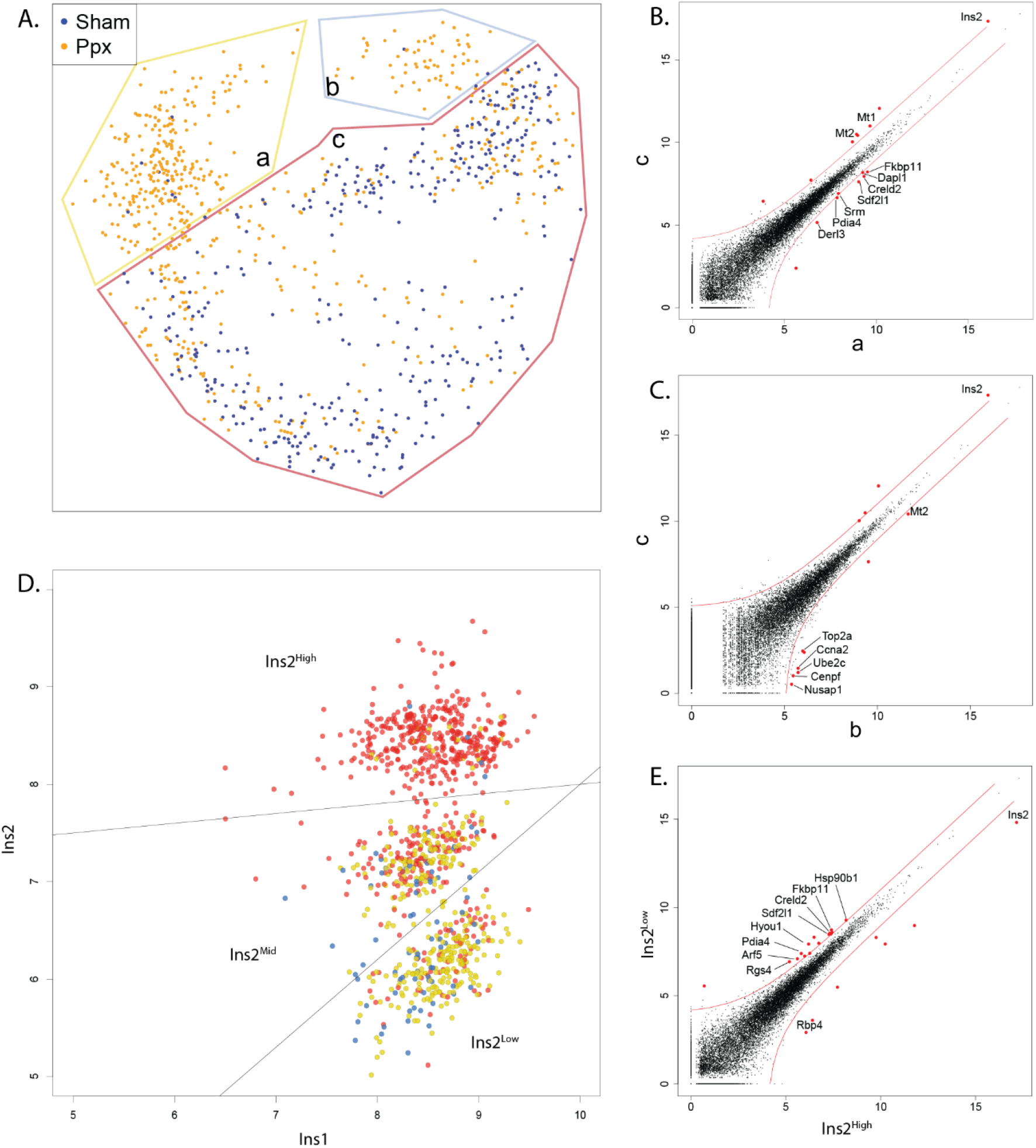
Heterogeneity of beta-cells in the regenerating endocrine pancreas. **(A)** Dimension reduction map depicting beta cell heterogeneity after regeneration with enrichment of two subsets (a, orange; b, blue) and a single subset that persists in basal conditions (c, red). Scatter plots of differential mRNA expression between beta-cell subsets highlights enrichment of stress-related mRNAs (**B**, subset a vs. c) and cell-division-cycle mRNAs (**C**, subset b vs. c) in regeneration. **(D)** Scatter-plot of Ins2 mRNA (y axis) versus Ins1 mRNA (x axis) in beta-cells reveals heterogeneity. Log2 of read counts. Cells colored according to clusters in Figure 4A. **(E)** Differential mRNA expression between beta-cells that express low and high levels of Ins2 mRNA. Unique molecular identifier (UMIs) normalized to total reads and to cell numbers.

Overall, our data reveal previously-overlooked cellular dynamics of the regenerating endocrine pancreas, including relative alpha-cell depletion, relative gamma-cell enrichment, considerable beta-cell heterogeneity and beta-delta pairs (Figure 3).

### Physically-interacting beta-delta cell pairs

We unexpectedly identified a relatively large number of beta-delta pairs and alpha-delta pairs that is untypical for MARS-seq data (Keren-Shaul et al., 2019); MARS-seq2.0). We sought to establish that these are indeed genuine doublets.

Because the typical number of transcripts is significantly larger than any single endocrine cell type, we excluded the possibility of a polyhormonal phenotype (Tukey’s multiple comparisons test of UMI distribution, p-value<<0.0001, Supplementary figure S2 A and B). In addition, because the transcriptome is predominated by beta-cell mRNAs, it is unlikely that data is gained from delta-cells contaminated by cell-free RNA from fragmented beta-cells.

We demonstrated that the pairing of delta to beta cells is specific and is significantly enriched, relative to other endocrine cell couples (randomization permutation test (R, v. 4.0.3), 10,000 iteration, p<0.0001). Since, beta-delta pairs are observed more than can be expected at random, it is unlikely to result from cell-cell adhesion in the tube after dissociation.

Then, we simulated *in silico* synthetic doublets, by summing the transcriptome of beta and delta cells (Supplementary figure S3 A and B). The gained synthetic transcriptome differs from real doublets. Thus, the expression of 21 mRNAs that are expressed > twofold in real pairs (Chi^2 statistics, p-value<0.01, (Supplementary Figure S4). mRNAs enriched in doublets more than expected include Dock3, a regulator of cytoskeletal organization and cell–cell interactions(Caspi and Rosin-Arbesfeld, 2008; Chen et al.,2009; Kashiwa et al., 2001) and several transcription factors of the zinc finger protein family (Ferguson et al., 2009) (Supplementary Figure S4). Overall, the study reveals genuine pairs of beta and delta cells in the endocrine pancreas.

### Delta-cells interact with particular beta-cell types

Several studies have described physical interactions of delta-cells with adjacent alpha or beta-cells in the tissue (Arrojo et al., 2019; Rorsman and Huising, 2018). We sought to interrogate the nature of physically interacting beta-delta conjugates, by an analytical pipeline for physically interacting cells, PIC-seq (Giladi et al., 2020). Evolution of 20,000 synthetic pairs within the PIC-seq QC process, reveals that they are conceivably similar to real pairs ((Giladi et al., 2020) Supplementary Figures S5 A-C).

PIC-seq characterized pure beta or delta cellular subsets (Figure 5A-E), and a joint signature of beta-delta pairs (Figure 5F), noting the relative contribution of beta- and delta-cell mRNAs to the superimposed profile was quantified (Figure 5G). Intriguingly, delta-cells consistently paired to one particular subset of beta cells, dubbed “int-beta” (interacting beta-cells). Enrichment of beta-delta pairs in the regenerating pancreas, proposes that such interactions may be beneficial in coping with the tissue or metabolic challenges, imposed by injury or regeneration.

**Figure 5.**
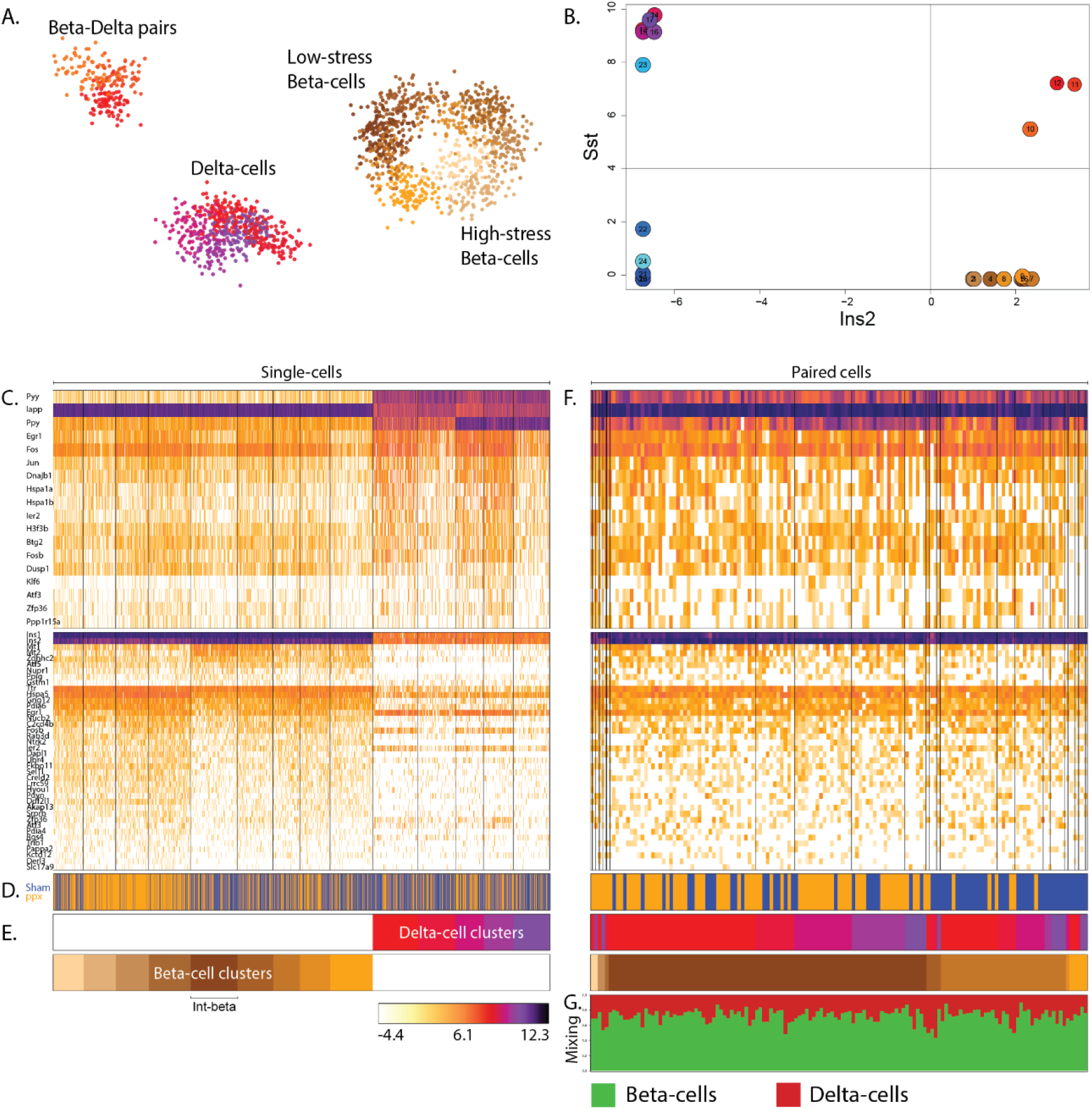
Physically interacting beta-delta cell pairs. **(A)** Two-dimensional projection of beta-cells, delta-cells, and beta-delta cell pairs. **(B)** Scatter-plot of 14 cellular subsets, demonstrating identification of pure beta or delta cell, by Ins2, or Sst and a joint signature with high expression of both hormones (subsets 10,11,12). **(C)** Heatmap of clustered RNA-sequencing data, 4 weeks after partial pancreatectomy defines 9 beta-cell subsets and 5 delta-cell subsets. Upper and lower panels indicate identity genes in delta- and beta-cell clusters, respectively. Data of 1500 cells collected from 6 pancreata analyzed by MetaCell (Baran et al., 2019). Sham-operated mice and mice lacking of lineage tracing included in the same analysis. Meta-cells (columns) and maximally enriched gene markers (rows). Color-bar indicators of **(D)** partitioning of cells derived from sham (blue) or ppx (orange) pancreata or **(E)** beta / delta meta-cells. Int-beta are interacting beta-cells. **(F)** Heatmap of clustered RNA-sequencing data, of 212 physically interacting beta-delta cell pairs, grouped by their contributing beta- and delta-cell identities. **(G)** Annotation and estimation of relative transcriptome contribution, derived from beta/delta cells (green/red).

### Beta-cells, juxtaposed to delta-cells, display a specific molecular signature

To validate the single-cell sequencing results, we orthogonally performed a single-molecule fluorescence *in situ* hybridization (smFISH) study for the detection of Somatostatin and Insulin2 (Figure 6 A-C). In addition, we studied stress-related Fkbp11 mRNA, which was suggested by the single cell sequencing to be one of the markers of regenerating beta-cells (Figure 4). The quantification revealed that beta-cells juxtaposed to delta-cells express less of the stress-related marker, Fkbp11 and more Ins2, relative to beta-cells distant from delta-cells in the regenerating pancreas, and relative to beta-cells that are paired to delta-cells under non-regeneration basal conditions (Figure 6 D, E, quantification of smFISH signal in 1361 cells, derived from 16 islets/ 6 pancreata. non-parametric Kolmogorov-Smirnov test). Together, two orthogonal approaches demonstrate that stress signature is dampened and Insulin expression is upregulated in beta-cells that are at physical interaction with delta-cells and that this is a molecular response that is observed specifically during regeneration.

**Figure 6.**
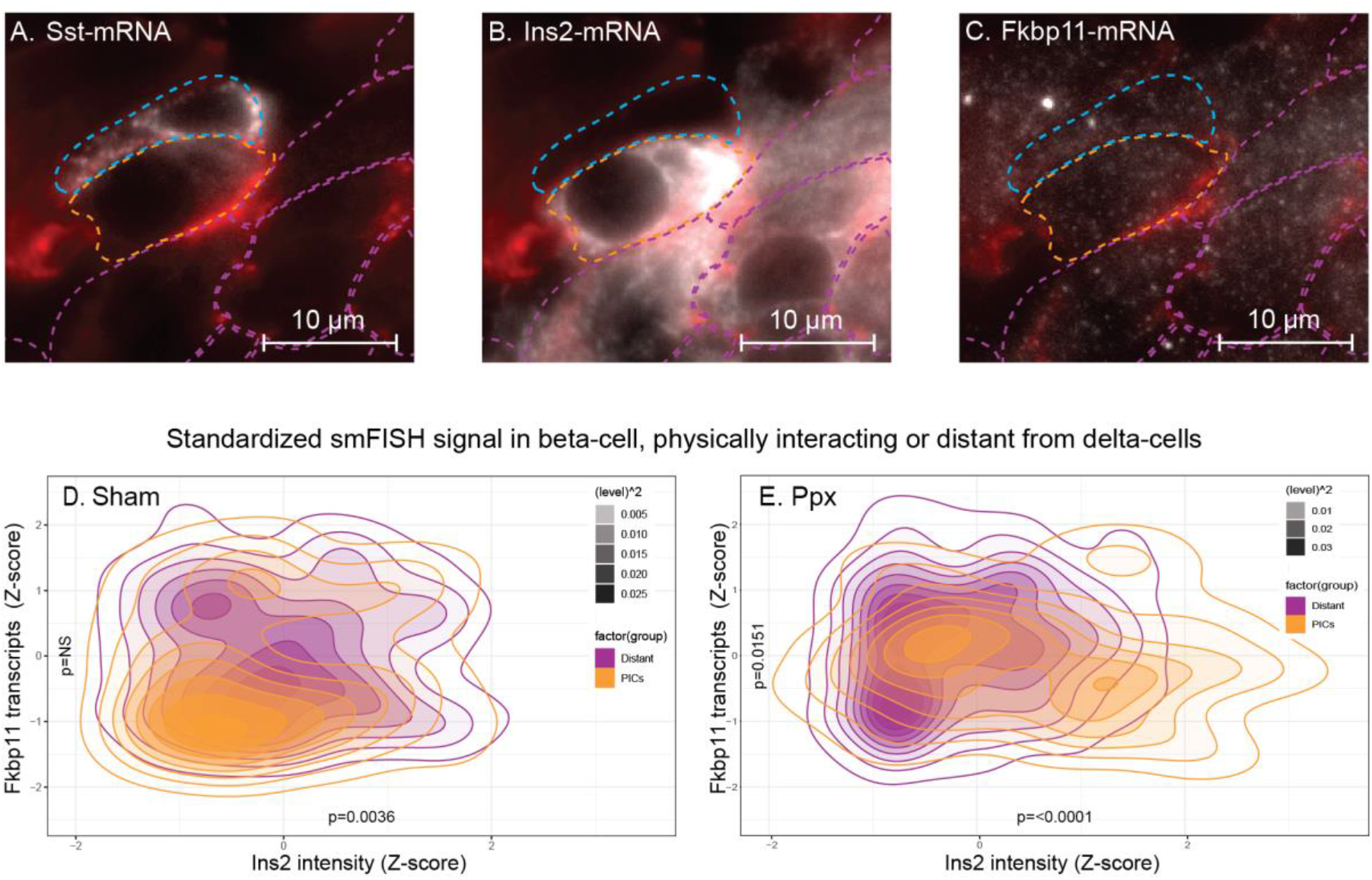
Molecular signature of physically interacting beta-delta cell pairs. smFISH study micrographs of Sst **(A)**, Ins2 **(B)** and Fkbp11 **(C)** mRNAs in islets of Langerhans. Beta-cells, juxtaposing delta-cells (orange), distant from delta-cells (purple), and delta-cells (blue). Cell membrane demonstrated by Phalloidin staining (red), smFISH signal (white). Density map of standardized Fkbp11 mRNA (x-axis) vs. Ins2 mRNA (y-axis) in 1361 cells, derived from 16 islets/ 6 pancreata. Beta-cells, juxtaposing delta-cells (orange, PIC) and beta-cells distant from delta-cells (purple), in islets from sham-operated mice (sham, **D**) or following pancreatectomy (Ppx, **E**). Data gained by quantification of smFISH signal in 1361 cells, derived from 16 islets/ 6 pancreata. P-values calculated by non-parametric Kolmogorov-Smirnov test (between distant vs. PICs for Fkbp11 and Ins2 transcript levels.

## Discussion

The emergence of single-cell transcriptome profiling paves the way to studies of endocrine pancreas heterogeneity at unprecedented cellular resolution. The current study reveals the cellular dynamics of the regenerating pancreas after surgical resection. It resonates some of the observations reporter by <u>Tatsuoka</u> et al (2020), were self-duplicating beta-cells were observed following partial pancreatectomy. Our work further enables a new look into beta-cell heterogeneity following regeneration and physically interacting endocrine cells, which were overlooked until the recent development of computational methods for discrimination of true interactions from sorting doublet artifacts.

We report that during regeneration there is a relative reduction in alpha-cell populations, accompanied by relative expansion of gamma-cells. In addition, beta-cell diversify. Thus, regeneration-associated beta-cell heterogeneity, appears to progress threefold: featuring ER-stress –associated markers (e.g., Fkbp11, Dapl1, Creld2), cell division cycle markers (e.g., Top2a, Ccna2, Ube2c) or beta-cell activity markers (high Ins2 along with Mt1, Mt2, ATF5 and Nupr1) accompanied with low ER stress markers.

We further report the unexpected dynamics in the delta-beta axis. We demonstrate that delta-cells of the regenerating pancreas can be denoted by the activation of a lineage tracer that is driven by the endogenous promoter of the transcription factor Sox-9. It is noticeable because in the mature organ Sox9 is primarily expressed in ductal and centero-acinar cells (Kopp et al., 2012; Seymour, 2014; Seymour et al., 2007). However, based on careful analysis we suggest that Sox9 descendant are most likely intrinsically endocrine, and we did not find any evidence that the source of the Sox-9 descendants might be of non-endocrine origin.

Delta cell population is expanding, approximately twofold, during regeneration. It inhabits the islet circumference in a zonated manner. These post-injury regeneration-related changes may be consistent with delta-cell hyperplasia in rodent diabetes models (Alan et al., 2015; Leiter et al., 1979).

Beta cells re-enter the cell division cycle in the regenerating pancreas, peaking at the first week after injury (Ackermann Misfeldt et al., 2008; Dor et al., 2004; Teta et al., 2007; Togashi et al., 2014) and were recently suggested to be associated with intricate activation of stress response, cell cycle progression effectors and tumor suppressors (Tatsuoka et al., 2020). Our analysis reveals a residual replicating beta cells, even 4 weeks following surgery, indicative of a prolonged replication phase than previously appreciated (Togashi et al., 2014).

We demonstrate an increase in the number of physically-pairing beta- and delta-cells in response to regeneration. This is in accordance with the increase in delta cell numbers. The distinctive molecular signature of beta-delta pairs in regeneration differs from beta-delta pairs under basal conditions, suggesting a regeneration-associated crosstalk. We demonstrate that only a particular subset, interacting beta cells (“Int-beta”), is capable of pairing to delta-cells. These beta cells display broad transcriptomic differences with reference to other beta cell subsets. However, it is unknown if the physical interaction with delta cell imposes broad transcriptional changes to beta-cells and / or that only specific subtypes of beta-cells are molecularly competent to induce beta-delta interactions.

Beta-delta proximity and crosstalk in the regenerating pancreas was described (Arrojo et al., 2019; Leiter et al., 1979), but could not be systematically quantified. Our study reveals that physical interaction with delta-cells is associated with reduced expression of stress markers and with augmented expression of mRNAs, typical of beta cell function. These two entities may be linked by the fact that high demand of insulin secretion imposes stress on beta-cells (Fonseca et al., 2011).

In summary we suggest that physically interacting delta-cells provide a unique protective niche that safeguards beta-cells from exhaustion. However, it is unknown if paracrine somatostatin or other means of cellular communication play a role in establishing the protective niche.

### Limitations

The study is first to perform analysis of cellular pairs in the endocrine pancreas. However, the functional importance of beta-delta crosstalk requires additional studies. Plausibly paracrine somatostatin plays a role in controlling beta-cell transcriptome and presumably function. However, non-secreted membrane-bound molecules might be additionally considered as means of communication. Our study focuses on a defined time point, 4 weeks after pancreatectomy, encouraging detailed future studies of the dynamics of beta-delta pairing in earlier phases of regeneration.

Delta cells extend compound filopodia-like protrusions to communicate with cells in their vicinity (Arrojo, 2019). Therefore, quantified pairing of beta and delta cells by microscopy and taking into account the potential of delta cells to affect beta cells that are considered by our calculation as ‘distant, non-interacting beta cells’, might underestimate the real effect of beta-delta pairing events.

In summary, the cellular dynamics of the regenerating endocrine pancreas are unfolded by single cell sequencing. Regeneration-associated heterogeneity of beta-cell, involves defined physical interactions with delta-cells that attenuate the load of stress and enable more robust beta cell function.

## Methods

### Animal experiments

Animal experiments were approved by the institutional Animal Care and Use Committee of the Weizmann Institute of Science. Mice were housed in a specific pathogen-free facility in individually ventilated cages on a strict 12-h light–dark cycle. C57BL/6 strain was purchased from Envigo. For lineage tracing, Sox9-Cre ^ERT2^ mice (Soeda et al., 2010) were crossed on to R26R-tdTomato conditional reporter [B6;129S6-Gt(ROSA)26Sortm9(CAG-tdTomato)Hze/J; (Madisen et al., 2010)]. 10 weeks old Sox9-Cre^ERT2^;tdTomato males were injected subcutaneously with tamoxifen (0.2-0.4 mg /g body weight 20-25 gr body weight, T5648, Sigma). Full clearance of residual tissue tamoxifen was allowed over 14 days, before any additional procedure. Subtotal surgical resection of the pancreas (partial pancreatectomy), was performed on isoflurane-anesthetized 12-week-old male mice, via midline abdominal incision, removing ~70%of the pancreas tissue, while preserving the pancreatic duct and splenic artery. Sham laparotomy and minimal rubbing of the pancreas with sterile q-tips served as control procedure. Abdominal muscles and skin were sutured using absorbable PGA 5/0 sutures (Intrag Medical Techs Co/ Ltd.). 0.1 mg/kg buprenorphine-HCl in normal saline was injected subcutaneously for analgesia, and enrofloxacin (2ml/400ml) given as prophylaxis oral antibiotic in drinking water for 7 days.

### Tissue microscopy

Islets of Langerhans were isolated by retrograde intra-ductal perfusion of pancreata with 5 ml 0.8 mg/ml Collagenase P following protocol described in (Szot et al., 2007). Freshly isolated islets of Langerhans or whole pancreata were fixed in 4% paraformaldehyde (PFA)/PBS at 4°C overnight, dehydrated in 30% sucrose, soaked in OCT (Tissue-Tek), frozen in mold on dry ice. Cryosections of 8 μm (Cryostat M3050S, Leica) were mounted onto Superfrost plus slides. For immunofluorescence, cryosections were airdried for 30 min, permeabilized with 0.2% Triton/TBS, incubated with Cas-block (008120 Thermo-fisher) for 10 min. and then incubated overnight at 4°C with primary antibodies, diluted in CAS-block. Sections were washed in PBS and incubated with secondary fluorescent antibody, mounted on glass slides with DAPI-mounting medium (Fluoroshield^™^ with DAPI, Sigma). Slides were air-dried over-night and examined with LSM 800 laser-scanning confocal microscope (Carl Zeiss), equipped with a Zeiss camera with ×20 / x40 or ×63 magnification (Thornwood, NY, USA). The following primary antibodies were used: guinea-pig anti Insulin (ab7842; Abcam, 1:200), goat anti-Sst (SC-7819; Santa Cruz biotechnology, 1:200) or rabbit anti-Sox9 (AB5535; Millipore, 1:200). All antibodies were previously validated, and immunostaining included negative controls (no primary antibody). Single Molecule FISH was performed as described in(Farack et al.,2018; Farack and Itzkovitz, 2020; Itzkovitz et al., 2011) using the Stellaris FISH Probe libraries (Biosearch Technologies, Inc., Petaluma, CA), coupled to Cy5 (GE Healthcare, PA25001), Alexa594 (Thermo Fisher, A37572) or TMR (Molecular Probes, C6123). (Table of probes TBD). Wash buffer and hybridization buffer contained 30% Formamide. Nuclei were counterstained with Dapi (Sigma-Aldrich, D9542) and cell borders, counterstained with alexa fluor 488 conjugated phalloidin (Thermo Fisher, A12379). Slides were mounted using ProLong Gold (Molecular Probes, P36934). Endocrine cells were detected by insulin or somatostatin signal. Micrographs captured by Nikon inverted fluorescence microscope eclipse ti2 series, equipped with a 100×oil-immersion objective and ixon ultra 888camera using NIS elements advanced research (Nikon). The image-plane pixel dimension was 0.13 μm. Quantification was performed on stacks of five optical sections, at 0.3 μm intervals, in which not more than a single cell was observed.

### Endocrine single-cell studies

Isolated Islets of Langerhans were dispersed to single-cell suspensions by incubation in a solution of 50% trypsin-EDTA (GIBCO) at 37°C for 3 min followed by gentile mechanic agitation, and stopped by adding 10% RPMI 1640 / FBS (vol/vol). For cell-sorting, islet cells were suspended in ice-cold sorting buffer (PBS supplemented with 0.2 mM ethylenediaminetetraacetic acid, pH 8 and 0.5% BSA), filtered through a 50-μm cell strainer and stained for viability by DAPI (1 ug/ml). Flow cytometry analysis and sorting were performed on a BD FACSAria Fusion instrument (BD Immunocytometry Systems), using a 100-μm nozzle, controlled by BD FACS Diva software v8.0.1 (BD Biosciences). Further analysis was performed using FlowJo software v10.2 (Tree Star). Either unstained, single stained tdTomato or DAPI only control cells were used for configuration and determining gates boundaries. tdTomato^+^ / DAPI ^−^ target cells were sorted into 384-well cell-capture plates containing 2 μl lysis solution and barcoded poly(T) reverse-transcription primers for single-cell RNA-seq. Empty wells in the 384-well plates served a no-cell controls. Immediately after sorting, each plate was spun down to ensure cell immersion into the lysis solution, and frozen at −80°C until processed. Single-cell libraries cDNA were prepared reverse-transcribed from messenger RNA of islet cells barcodes (Jaitin et al., 2014; Keren-Shaul et al., 2019). ScRNA-seq libraries were pooled at equimolar concentrations and sequenced on an Illumina NextSeq 500 at a median sequencing depth of 24,500 reads per cell. Sequences were mapped to the mouse genome (mm10) using HISAT2 (Kim et al., 2019), demultiplexed and filtered to exclude reads outside exons or with multiple mapping positions. Cell libraries with 500 - 500,000 UMIs and <20% mitochondrial mRNAs were included in downstream MetaCell analysis (Baran et al., 2019), which derives cohesive groups of cellular profiles. To study physical-interaction between cells, we employed PIC-seq (Giladi et al., 2020) on single-cell data.

## Supplementary figures

**Supplementary Figure 1A.**
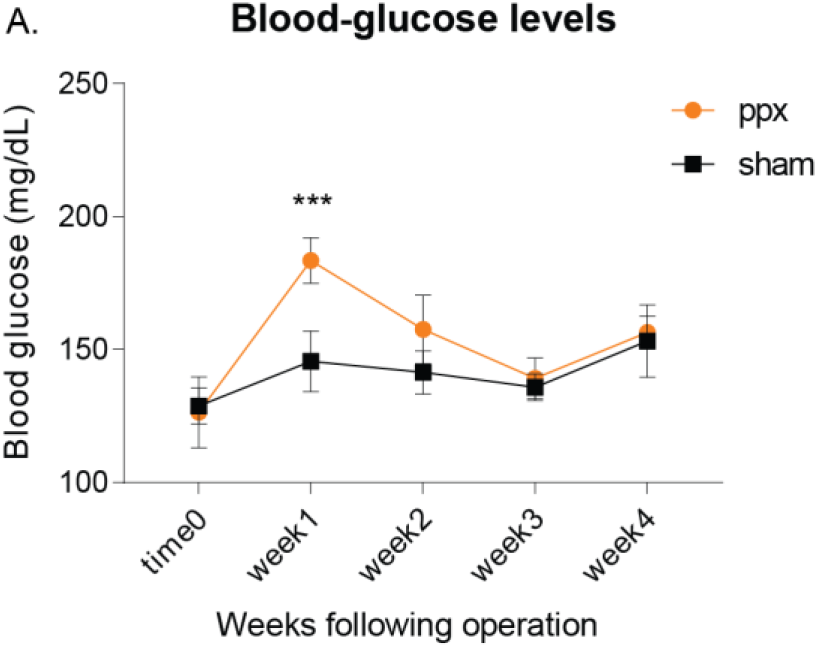
Blood glucose levels in partially-pancreatectomized (orange) or sham-operated mice (black), in the course of four weeks, following surgical procedure. 2way anova, Sidak’s multiple comparisons test, *** p-value = 0.0001. Data obtained from 5 mice from 2 experimental groups.

**Supplementary Figure 2.**
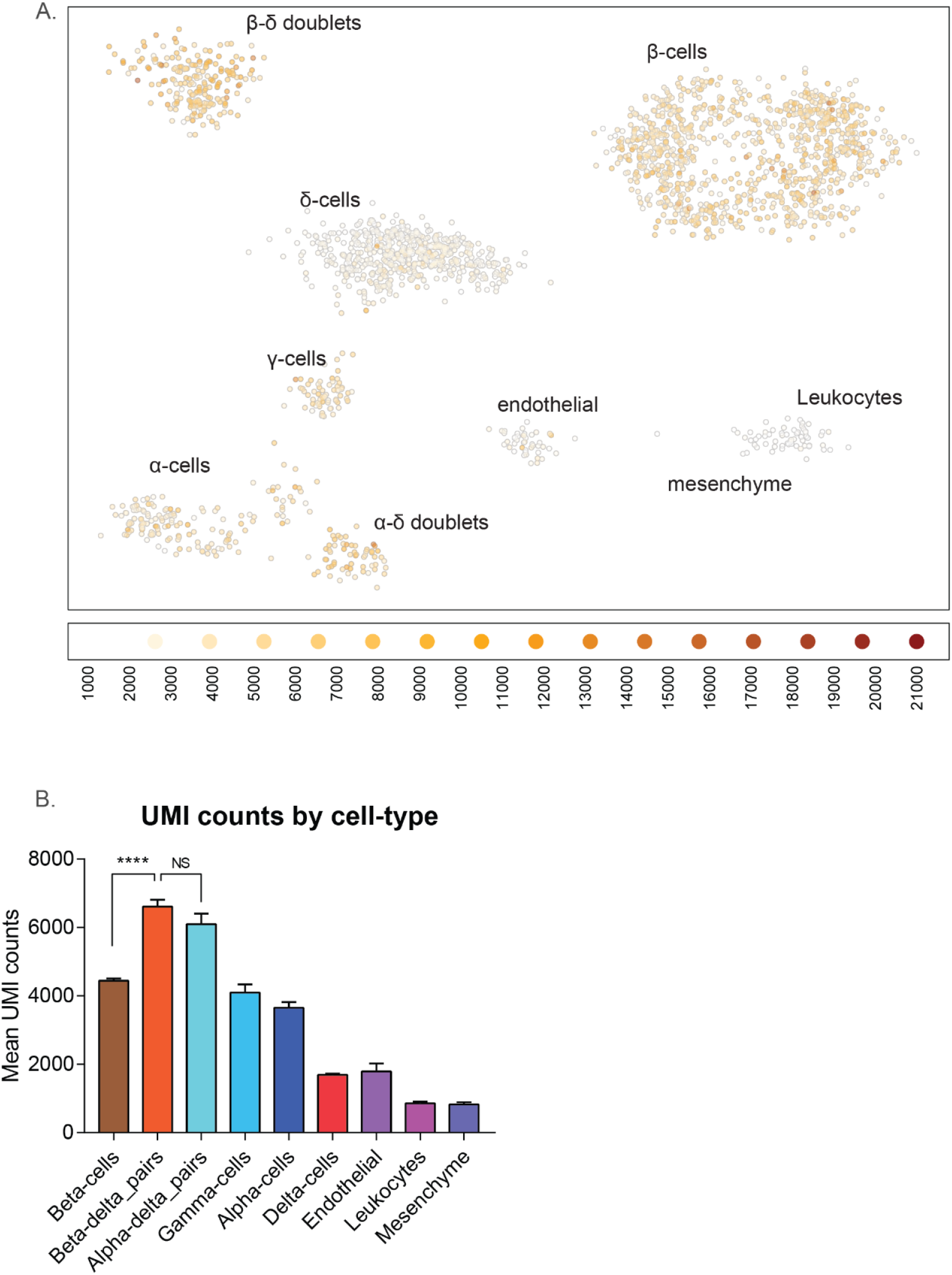
**(A).** Dimensional reduction map from MetaCell analysis, illustrating UMI counts of single cells across MetaCells. **(B)** Bar-plot of cell-type specific mean UMI-counts. Data of 2104 cells following MetaCell analysis. Tukey’s multiple comparisons test. **** p-value <0.0001. NS = non-significant. Cell-types were inferred according to unique transcripts. 974 beta-cells, 207 beta-delta pairs, 58 alpha-delta pairs, 66 gamma-cells, 151 alpha-cells, 539 delta-cells, 46 Endothelial cells, 54 Leukocytes, and 9 mesenchyme cells.

**Supplementary Figure 3.**
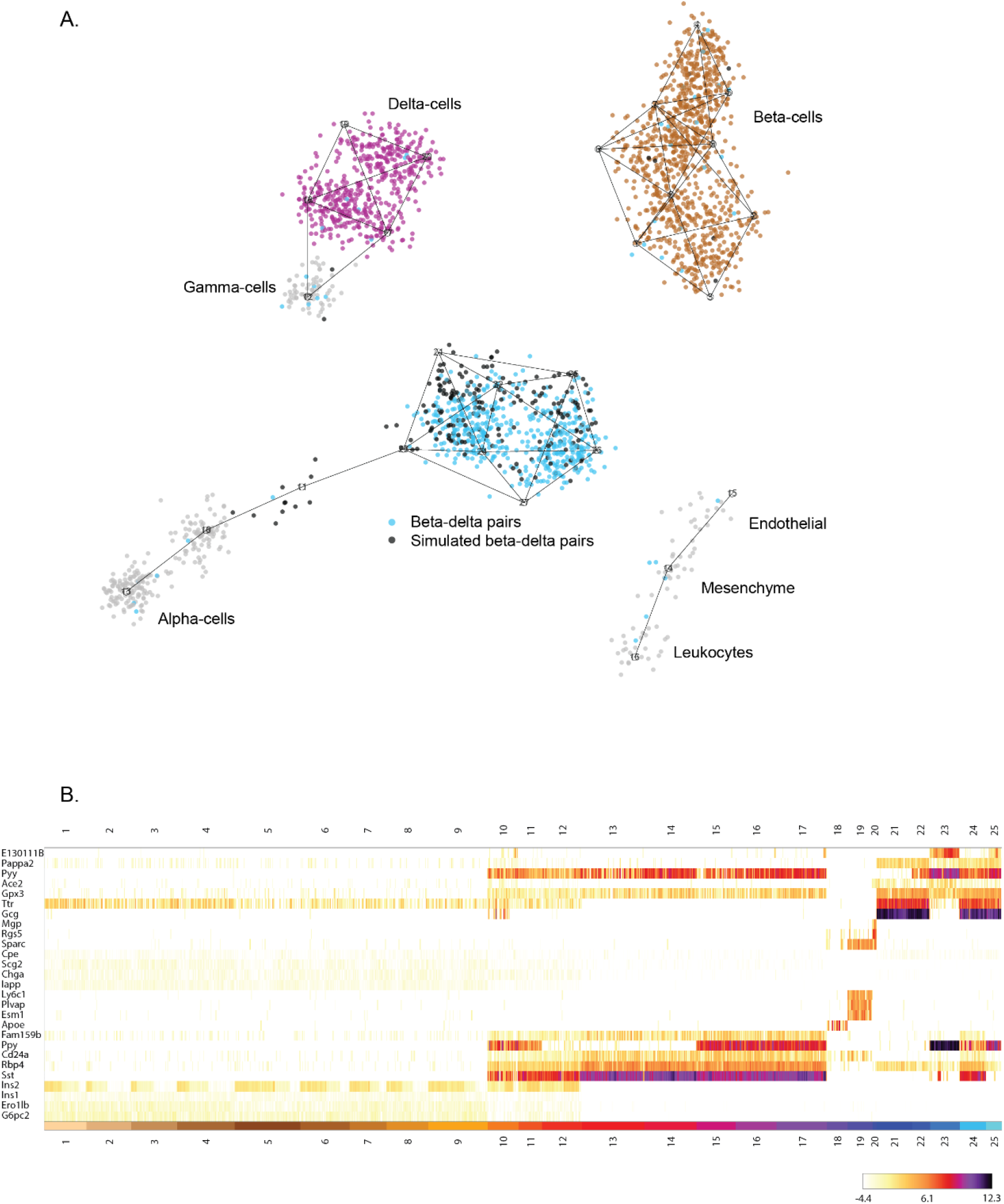
**(A).** Dimensional reduction map from MetaCell analysis, illustrating simulated doublets (black dots) from single beta-(brown dots) and delta-cells (purple dots). Blue dots represent beta-delta cell pairs. **(B)** Heatmap of clustered RNA-sequencing data from Sox9 lineage-traced islet cells, 4 weeks following partial pancreatectomy. Data of 2300 cells collected from 6 pancreata analyzed by MetaCell. Data includes 515 simulated beta-delta cells.

**Supplementary Figure 4.**
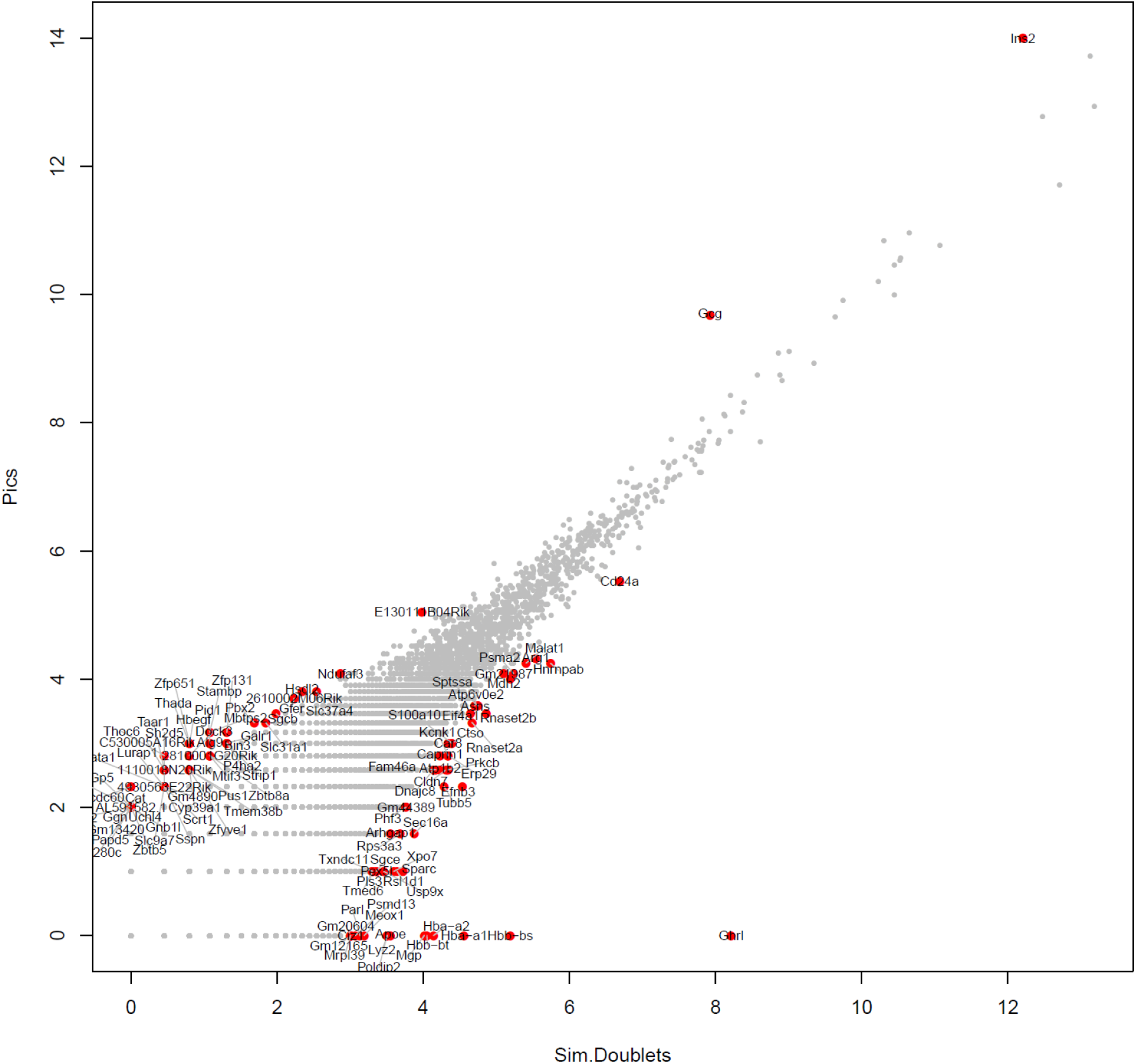
Scatter plots of differential mRNA expression between simulated beta-delta pairs (x axis) and observed beta-delta pairs (y-axis) highlights enrichment of genes supporting beta-cell function and cellular adhesion molecules.

**Supplementary Figure 5.**
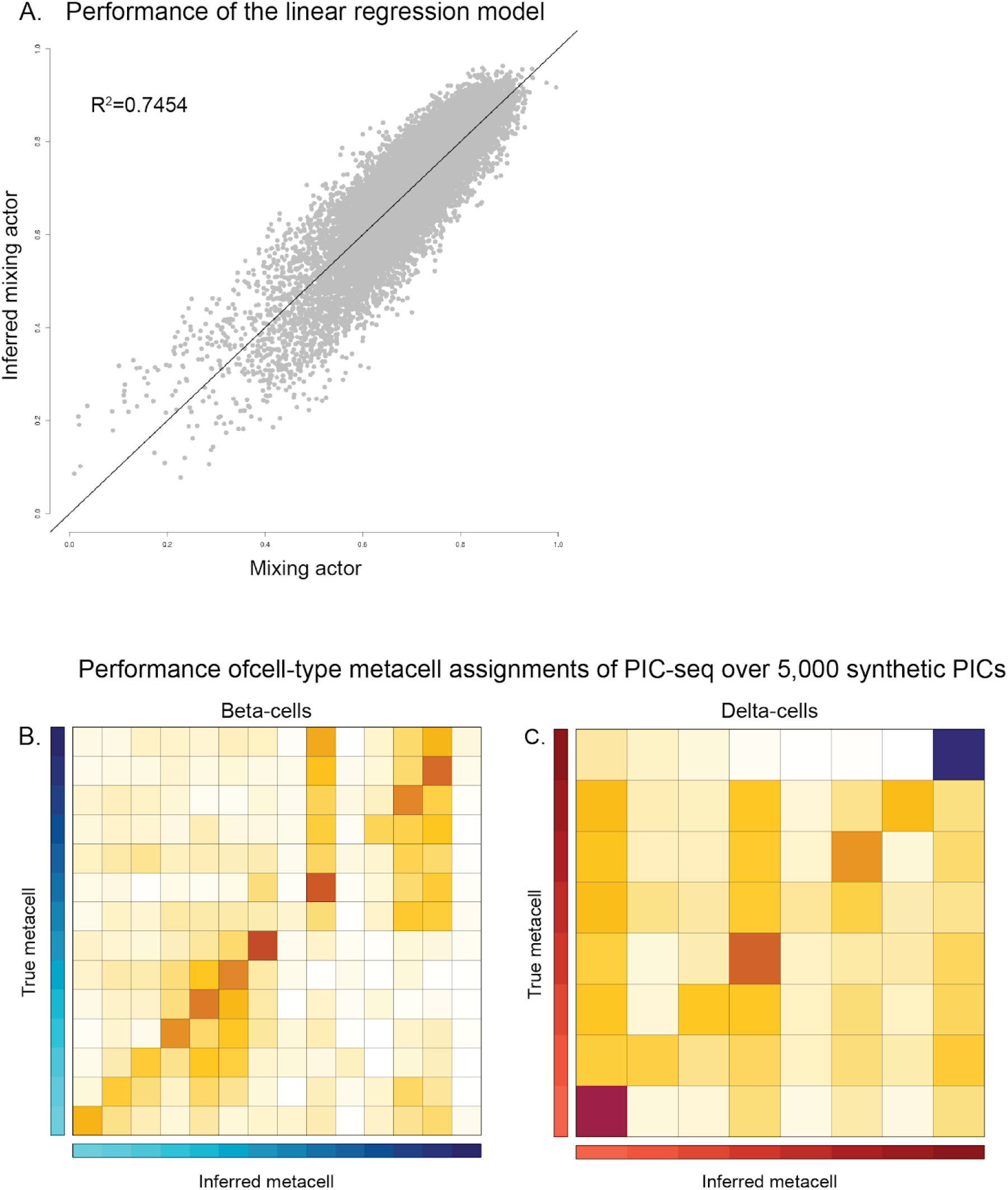
**(A)**. Performance of the linear regression model, used to estimate the mixing factor of 20,000 synthetic PICs. **(B, C)** Performance of the **(B)** beta-cell and **(C)** Delta-cell MetaCell assignments of PIC-seq over 5,000 synthetic PICs. Each row summarizes all synthetic PICs originating from one MetaCell and their assignments to MetaCell by PIC-seq (columns. Data is row-normalized).

## Acknowledgments

E.H. is the Mondry Family Professorial Chair and head of the Nella and Leon Benoziyo Center for Neurological Diseases at Weizmann Institute of Science. We thank members of the Hornstein lab for helpful critiques. We would like to thank Dr. Tomer-Meir Salame, Dana Hirsch and Dr. Liat Fellus-Alyagor, from the Life Sciences Core Facilities, Weizmann Institute of Science, for technical support. The work was funded by European Research Council under the European Union’s Seventh Framework Programme (FP7/2007- 2013)/ERC grant agreement 617351 and the Israel Science Foundation (135/16); Research in the Hornstein lab is further supported by Radala Foundation; Minerva Foundation with funding from the Federal German Ministry for Education and Research, ISF Legacy grant 828/17; Yeda-Sela, Yeda-CEO; Benoziyo Center Neurological Disease; Weizmann - Brazil Center for Research on Neurodegeneration at The Weizmann Institute of Science, Vener New Scientist Fund; Julius and Ray Charlestein Foundation; Fraida Foundation; Wolfson Family Charitable Trust; Abney Foundation; Merck; Maria Halphen and the estates of Fannie Sherr, Lola Asseof, Lilly Fulop, and E. and J. Moravitz and by Dr. Sydney Brenner and friends.

